# The global distribution and spread of the mobilized colistin resistance gene *mcr-1*

**DOI:** 10.1101/220921

**Authors:** Ruobing Wang, Lucy van Dorp, Liam Shaw, Phelim Bradley, Qi Wang, Xiaojuan Wang, Longyang Jin, Qing Zhang, Yuqing Liu, Adrien Rieux, Thamarai Dorai-Schneiders, Lucy Anne Weinert, Zamin Iqbal, Xavier Didelot, Hui Wang, Francois Balloux

**Affiliations:** Department of Clinical Laboratory, Peking University People’s Hospital, Beijing, 100044, China; UCL Genetics Institute, University College London, Gower Street, London WC1E 6BT, UK; Wellcome Trust Centre for Human Genetics, University of Oxford, Oxford OX3 7BN, UK; Institute of Animal Science and Veterinary Medicine, Shandong Academy of Agricultural Sciences, Shandong Province, Jinan, 250100, China; UMR PVBMT, CIRAD, 97410 St Pierre, Reunion, France; Division of Infection and Pathway Medicine, 49 Little France Crescent, Edinburgh EH16 4SB, UK; Department of Veterinary Medicine, Cambridge CB3 0ES, UK; Department of Infectious Disease Epidemiology, Imperial College London, Norfolk Place, 21 London W2 1PG, UK; European Molecular Biology Laboratory, European Bioinformatics Institute (EMBL-EBI), Wellcome Genome Campus, Cambridge CB10 1SD, UK

**Author notes:** Correspondence: Hui Wang and Francois Balloux.

## Abstract

Colistin represents one of the very few available drugs for treating infections caused by carbapenem resistant Enterobacteriaceae (CRE). As such, the recent plasmid-mediated spread of the mobilized colistin resistance gene *mcr-1* poses a significant public health threat requiring global monitoring and surveillance. In this work, we characterize the global distribution of *mcr-1* using a dataset of 457 *mcr-1* positive sequenced isolates consisting of currently publicly available *mcr-1* carrying sequences combined with an additional 110 newly sequenced *mcr-1* positive isolates from China. We find *mcr-1* in a diversity of plasmid backgrounds but identify an immediate background common to all *mcr-1* sequences. Our analyses establish that all *mcr-1* elements in circulation descend from the same initial mobilization of *mcr-1* by an ISA*pl1* transposon in the mid 2000s (2002-2008; 95% higher posterior density), followed by a dramatic demographic expansion, which led to its current global distribution. Our results provide the first systematic phylogenetic analysis of the origin and spread of *mcr-1*, and emphasize the importance of understanding the movement of mobile elements carrying antibiotic resistance genes across multiple levels of genomic organization.

## Introduction

Colistin was largely abandoned as a treatment for bacterial infections in the 1970s due to its high toxicity and low renal clearance, but has been reintroduced in recent years as an antibiotic of ‘last resort’ against multi-drug-resistant (MDR) infections^1^. It is therefore alarming that the prevalence of resistance to colistin has become a significant concern, following the identification of plasmid-mediated colistin resistance conferred by the *mcr-1* gene in late 2015^2^. Colistin resistance is emblematic of the growing problems of antimicrobial resistance worldwide.

Up until 2015, resistance to colistin had only been linked to mutational and regulatory changes mediated by chromosomal genes ^3,4^. The mobilized colistin gene *mcr-1* was first described in a plasmid carried by an *Escherichia coli* isolated in China in April 2011^2^. The presence of colistin resistance on mobile genetic elements poses a significant public health risk, as these can spread rapidly by horizontal transfer, and may entail a lower fitness cost^5^. At the time of writing, *mcr-1* has been identified in numerous countries across five continents. Significantly, *mcr-1* has also been observed on plasmids containing other antimicrobial resistance genes such as carbapenemases^6^^–^^8^ and extended-spectrum β-lactamases (ESBL)^9^^–^^11^.

The *mcr-1* element has been characterized in a variety of genomic backgrounds, consistent with the gene being mobilized by a transposon^12^^–^^16^. To date, *mcr-1* has been observed on a wide variety of plasmid types, including IncI2, IncHI2, and IncX4^15^. Intensive screening efforts for *mcr-1* have found it to be highly prevalent in a number of environmental settings, including the Haihe River in China^17^, recreational water at public urban beaches in Brazil^18^, and faecal samples from otherwise healthy individuals^19,20^. While both Brazil and China have now banned the use of colistin in agriculture, the evidence that *mcr-1* can spread within hospital environments even in the absence of colistin use^19^ as well as in the community^20^ raises the possibility that the spread of *mcr-1* will not be contained by these bans.

The global distribution of *mcr-1* over at least five continents is well documented, but little is known about its origin, acquisition, emergence, and spread. In this study, we aim to shed light on these fundamental issues using whole genome sequencing data from 110 novel *mcr-1* positive isolates from China in conjunction with an extensive collection of publicly available sequence data sourced from the NCBI repository as well as the Short Read Archive (SRA).

Our data and analyses support an initial single mobilization event of *mcr-1* by an ISA*pl1-mcr-1-orf*-ISA*pl1* transposon around 2006. The transposon was immobilized on several plasmid backgrounds following the loss of the flanking ISA*pl1* elements, and spread through plasmid transfer. The current distribution of *mcr-1* points to a possible origin in Chinese livestock. Our results illustrate the complex dynamics of antibiotic resistance genes across multiple embedded genetic levels (transposons, plasmids, bacterial lineages and bacterial species), previously described as a nested “Russian doll” model of genetic mobility^21^.

## Results

### Dataset

We compiled a global dataset of 457 *mcr-1* positive isolates (Figure 1A), including 110 new whole genome sequences (WGS) from China, of which 107 were sequenced with Illumina short reads and three with PacBio long read technology. 195 isolates were sourced from publicly available assemblies in the NCBI sequence repository (73 completed plasmids, 1 complete chromosome, 121 assemblies). A further 153 sequences were sourced from the Short Read Archive (NCBI-SRA), after being identified as *mcr-1* positive using a k-mer index of a snapshot of the SRA as of December 2016 (see Methods). The whole dataset consists of 256 short-read datasets, 6 long-read PacBio WGS, 121 draft assemblies, and 74 completed assemblies.

**Figure 1.**
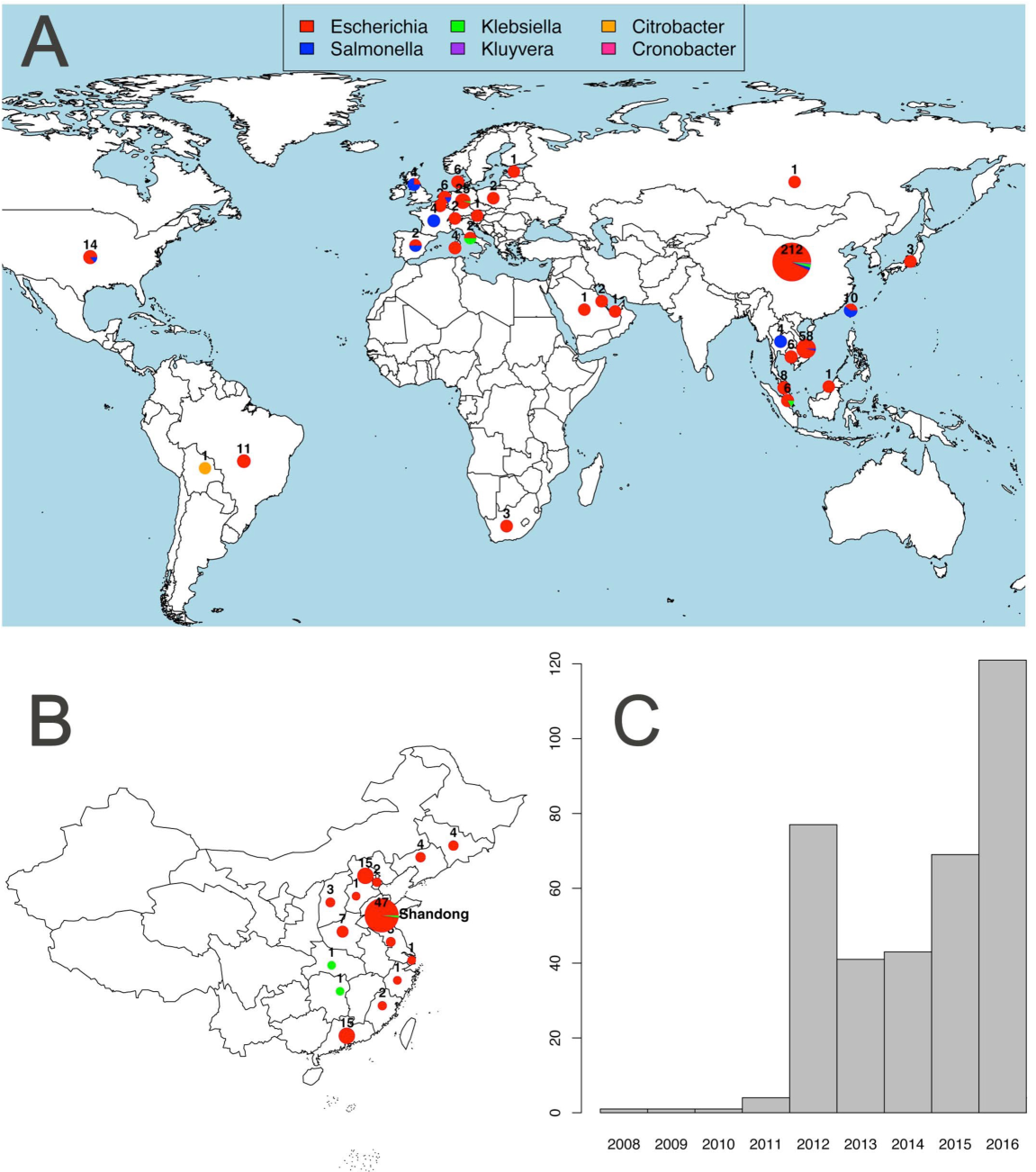
Overview of the *mcr-1* positive isolates included. **A**. Global map of *mcr-1* positive isolates included coloured by genus with the number and size of pies providing the sample size per location; **B**. Map of novel Chinese isolates sequenced for this study; **C**. Histogram of sampling dates (years) of the isolates.

Isolates carrying *mcr-1* were identified from 31 countries (Figure 1A). The countries with the largest numbers of *mcr-1* positive samples are China (212), Vietnam (58) and Germany (25). Within China, nearly half (45%) of positive isolates stem from Shandong province (Figure 1B). The vast majority of *mcr-1* positive isolates belong to *Escherichia coli* (411), but the dataset also comprises *mcr-1* positive isolates from another seven bacterial species: *Salmonella enterica* (29), *Klebsiella pneumoniae* (8), *Escherichia fergusonii* (2), *Kluyvera ascorbata* (2), *Citrobacter braakii* (2), *Cronobacter sakazakii* (1) and *Klebsiella aerogenes* (1) (Figure 1A). The majority of isolates for which sampling dates were available (80%), were collected between 2012 and 2016, with the oldest available isolates dating back to 2008 (Figure 1C).

The large number of *mcr-1* positive isolates from China, and the high incidence in Shandong province can be largely ascribed to the inclusion of our 110 newly sequenced isolates including 49 from Shandong and to another 37 isolates from a previous large sequencing effort^13^. However, even after discounting the isolates from these two sources, China remains, together with Vietnam one of the two countries with the highest number of sequenced *mcr-1* positive isolates.

### Evolutionary model

It has been proposed that *mcr-1* is mobilized by a composite transposon formed of a ~2,600bp region containing *mcr-1* (1,626bp) and a putative open reading frame encoding a PAP2 superfamily protein (765bp), flanked by two ISA*pl1* insertion sequences^12^. ISA*pl1* is a member of the IS30 family of insertion sequences, which utilize a ‘copy-out, paste-in’ mechanism with a targeted transposition pathway requiring the formation of a synaptic complex between an inverted repeat (IR) in the transposon circle and an IR-like sequence in the target. Snesrud and colleagues^12^ hypothesized that after the initial formation of such a composite transposon, these insertion sequences would have been lost over time, leading to the stabilization of *mcr-1* in a diverse range of plasmid backgrounds (Figure 2). In the following, we sought to test this model by performing an explicit phylogenetic analysis of the region surrounding *mcr-1* using our comprehensive global dataset.

**Figure 2.**
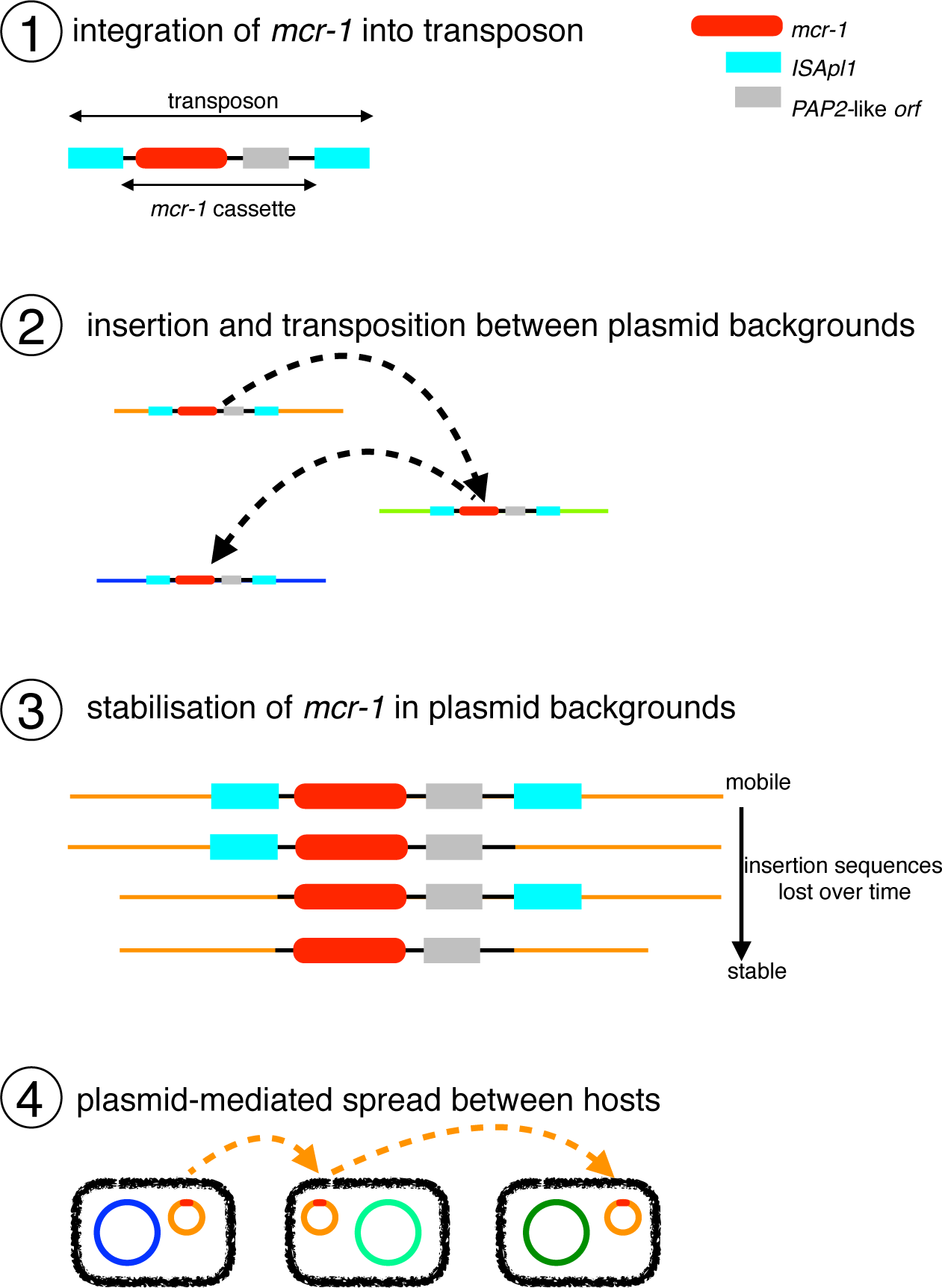
Schematic representation of the evolutionary model for the steps in the spread of the *mcr-1* gene. (1) The formation of the original composite transposon, followed by (2) transposition between plasmid backgrounds and (3) stabilisation via loss of ISA*pl1* elements before (4) plasmid-mediated spread.

### *Immediate genomic background of* mcr-1

If there had been a unique formation event for the composite transposon, followed by progressive transposition and loss of insertion sequences, we would expect to be able to identify a common immediate background region for *mcr-1* in all samples. Indeed, we were able to identify and align a shared region or remnants of it in all 457 sequences surrounding *mcr-1* (see Methods), supporting a single common origin for all *mcr-1* elements sequenced to date (Figure 3a). The majority of the sequences contained no trace of ISA*pl1* (n=260) indicating that the *mcr-1* transposon had been completely stabilized in their genomic background. 42 sequences contained indication of the presence of *ISApl1* both upstream and downstream, either in full copies (n=16), a full copy upstream and a partial copy downstream (n=7), a partial copy upstream and a full copy downstream (n=1), or partial copies upstream and downstream (n=18). Some sequences only had ISA*pl1* present upstream as a complete (n=55) or partial (n=99) sequence, and one sequence had only a partial downstream ISA*pl1* element. The downstream copy of ISA*pl1* was inverted in some sequences (n=3) and some sequences had full copies of ISA*pl1* present elsewhere on the same contig (n=7), consistent with its high observed activity in transposition^22^.

**Figure 3.**
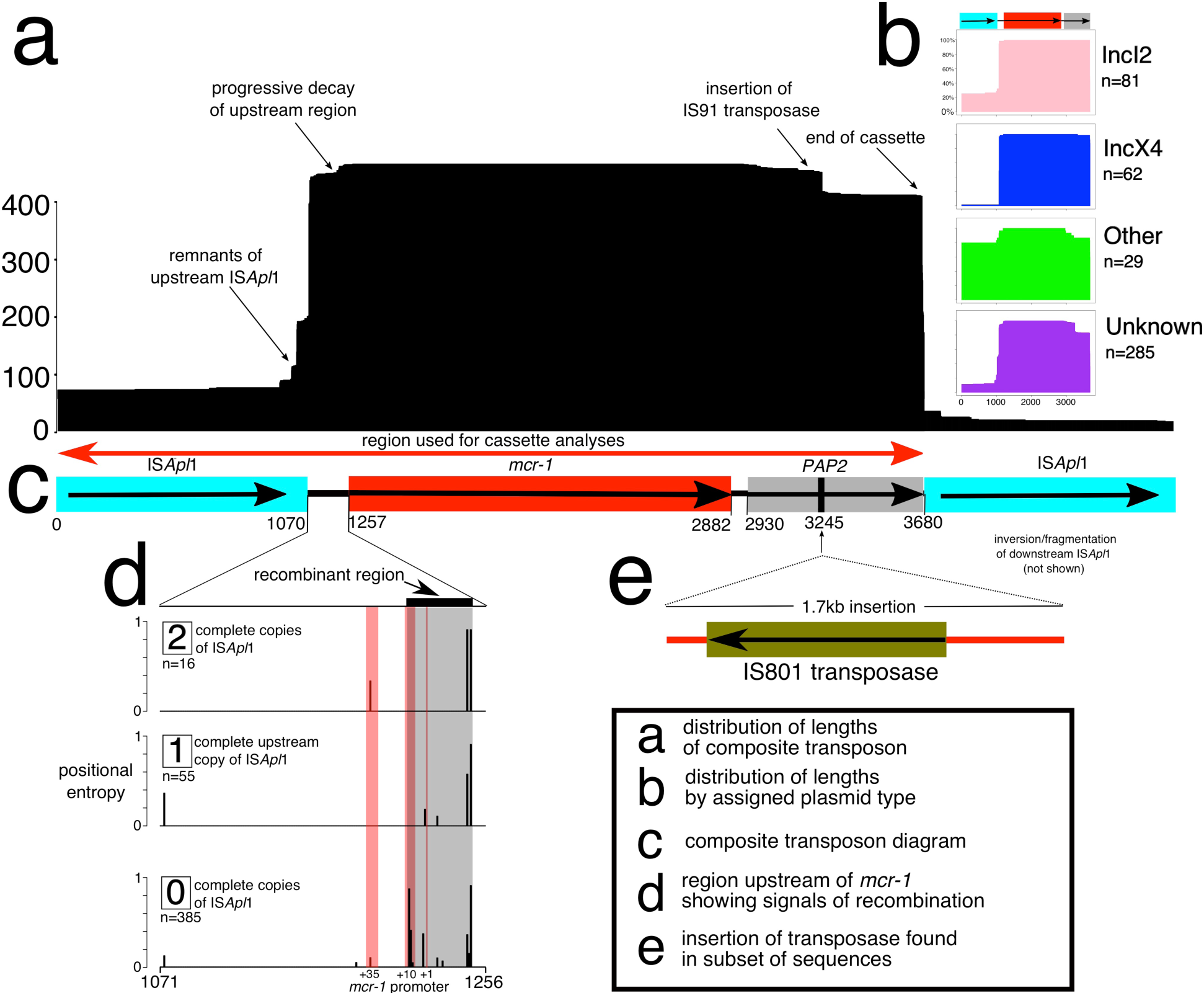
The genetic element carrying *mcr-1* is a composite transposon and is alignable across our global dataset. (a) Length distribution of the alignment across sequences. (b) Length distribution subset by plasmid type. (c) The composite transposon, consisting of IS*Apl*1, *mcr-1*, a PAP2 *orf*, and ISA*pl1*. The region indicated by the red arrow was used in phylogenetic analyses, after the removal of recombination. (d) The 186bp region upstream of *mcr-1* showed strong signals of recombination (grey box) that coincided with the promotor regions of *mcr-1* (red box), and this diverse region was removed from the subsequent alignment. (e) 28 sequences from Vietnam had a 1.7kb insertion containing a region with >99% sequence identity to IS801 transpose, suggesting subsequent rearrangement after initial mobilization.

Further inspection of the transposon alignment revealed that the 186bp region between the 3’ end of the upstream ISA*pl1* and *mcr-1* contained IR-like sequences similar to the IRR and IRL of ISA*pl1* (respectively: 93-142bp, 23/50 identity; and 125-175bp, 21/50 identity). The most variable positions in this 186bp region were at 177bp and 142bp, approximately coinciding with the end of the alignment with the IRs and were more variable in sequences lacking *ISApl1*, suggesting possible loss of function of the transposition pathway associated with *ISApl1* (Figure 3d). Some of these SNPs occurred in a stretch previously identified as the promoter region for *mcr-1^23^*, and this region showed strong signals of recombination. A small number of sequences (3%) had SNPs present in *mcr-1* itself. These tended to be at the upstream/5’ end of the sequence, particularly in the first three positions. A subset of the sequences from Vietnam (n=28) included a secondary 1.7kb insertion downstream of *mcr-1* containing an \S801 transposase, indicating subsequent rearrangements involving this region after initial mobilization of the transposon (Figure 3e).

To reconstruct the phylogenetic history of the composite *mcr-1* transposon, we created a sequence alignment for 457 sequences (Figure 3c) after removal of recombinant regions identified with ClonalFrameML, including the region immediately upstream of *mcr-1* between positions 1,212-1,247 (Figure 3d). A midpoint-rooted maximum parsimony phylogeny showed that there was a dominant sequence type with subsequent diversification, likely indicating the ancestral form of the composite transposon (Figure 4).

**Figure 4.**
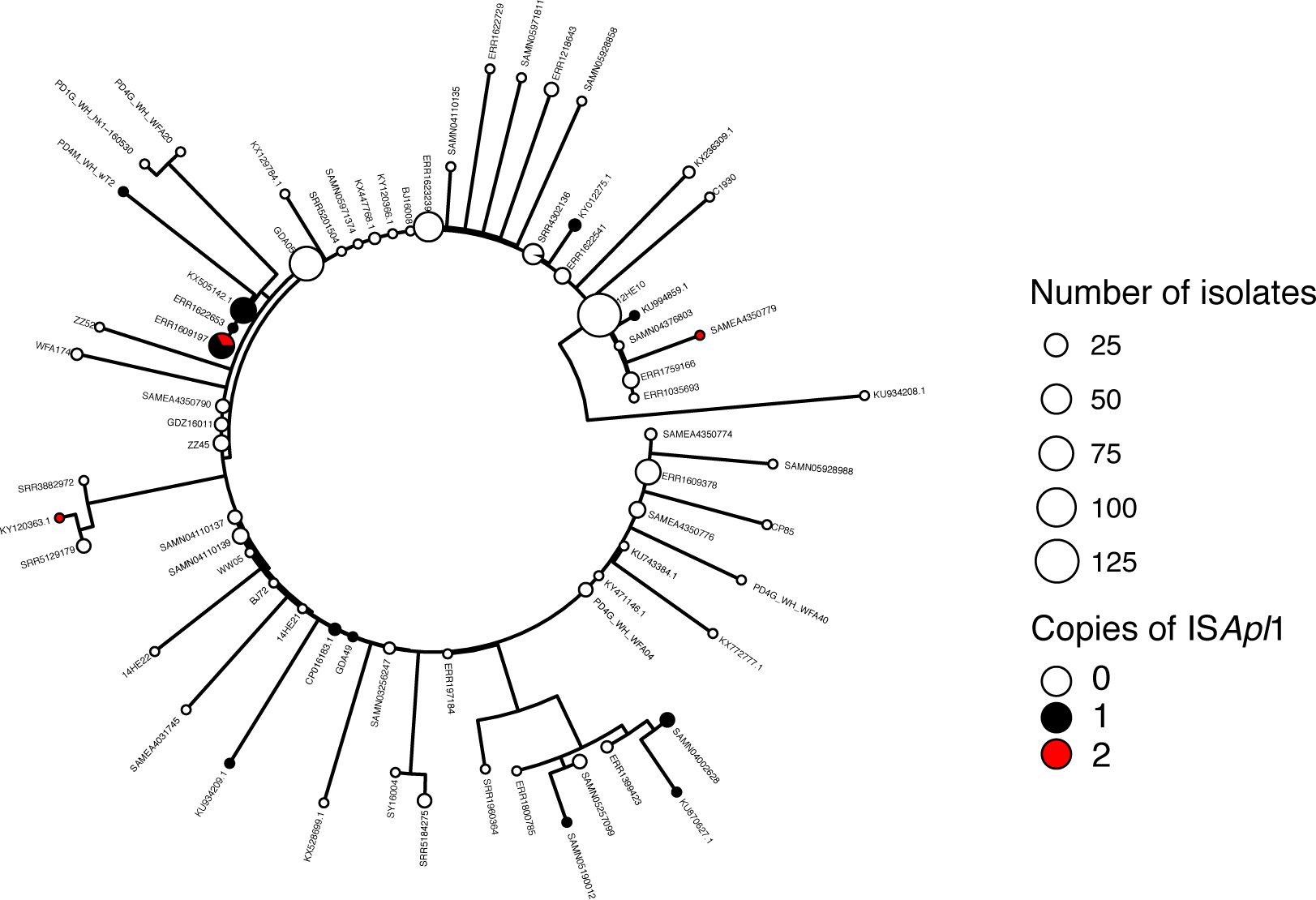
Phylogeny of the *mcr-1* composite transposon indicates a dominant sequence type with subsequent diversification. Midpoint-rooted maximum parsimony phylogeny based on the 3,522bp alignment of 457 sequences (recombinant regions removed). Size of points indicates the number of identical sequences, with a representative sequence for each shown next to each tip.

A Bayesian dating approach (BEAST) was applied to infer a timed phylogeny of the maximal alignable region of the *mcr-1* carrying transposon (see Methods). Based on this 3,522 site alignment we infer a common ancestor for 364 dated isolates in 2006 (Figure S1; 2002-2008 95% HPD strict clock, coalescent model) with a mutation rate around 7.51×10^−5^ substitutions per site per year (Table S1). There was no clear overall geographic clustering in the maximum clade credibility tree (Figure S2).

### *Wider genomic background of* mcr-1

Next, we explored the wider genomic background upstream and downstream of the conserved transposon sequences. We had sufficiently long assembled contigs for 172 isolates to identify plasmid types based on co-occurrence with plasmid replicons (see Methods) and identified *mcr-1* in 13 different plasmid backgrounds. IncI2 and IncX4 were the dominant plasmid types, accounting for 47% and 36% of the isolates, respectively (Figure 5). One isolate in our dataset was located on a chromosome.

**Figure 5.**
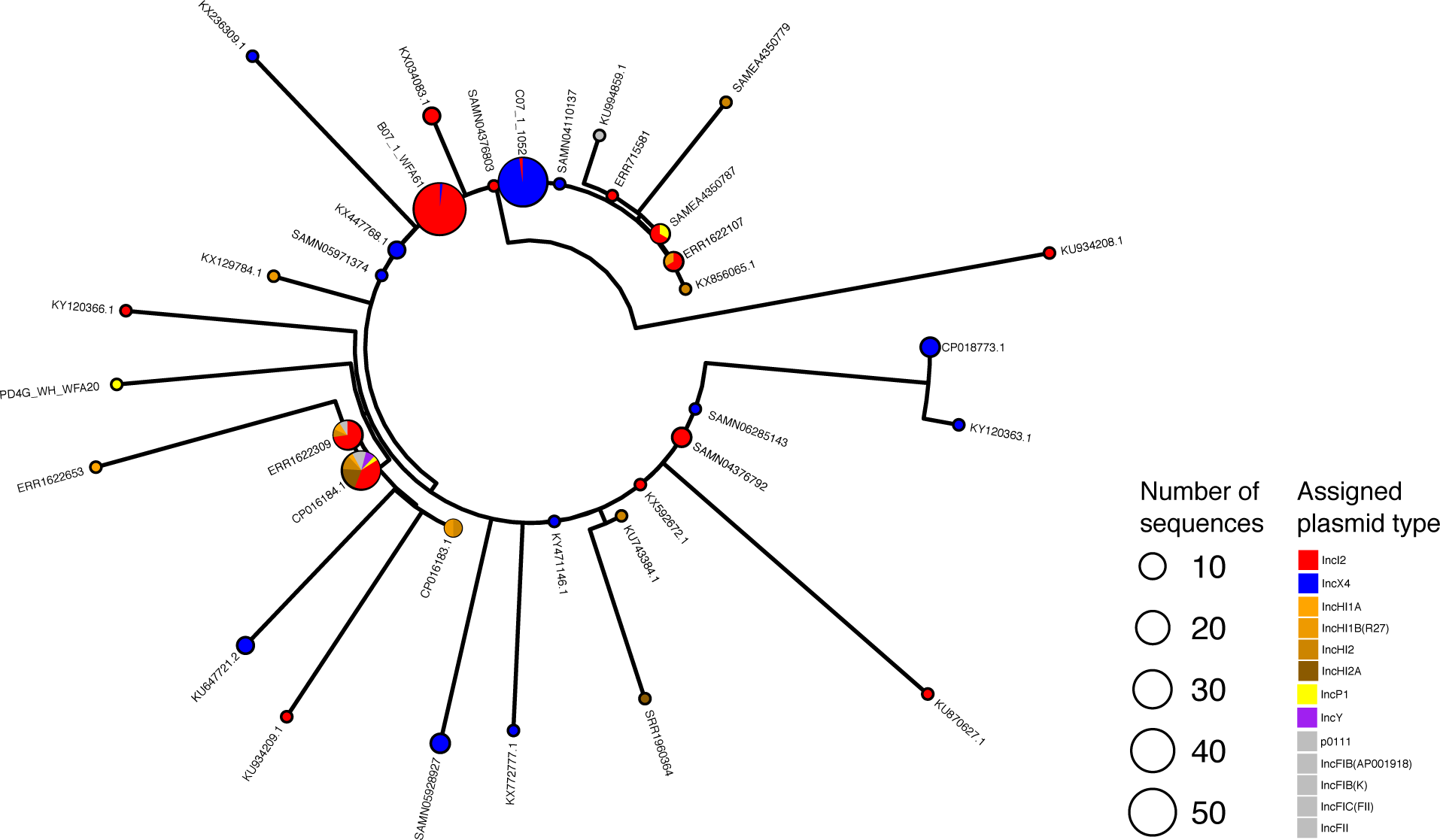
The distribution of plasmid types shown on the transposon phylogeny. Maximum parsimony tree (homoplastic sites removed, mid-point rooted, as in Figure 4) based on the composite transposon alignment for 172 sequences containing a plasmid replicon on the same contig i.e. those with an assigned plasmid type (color). IncI2 and IncX4 are the most common plasmid types. An example sequence ID is shown for each unique sequence.

The distribution of transposons carrying one or two copies of ISA*pl1* was highly heterogeneous across these plasmid types. For example, sequences with one or two copies of ISA*pl1* were found on 11 and seven types, respectively, which supports their mobility compared to those without ISA*pl1*, which were found in six plasmid types. Of the contigs carrying one copy of ISA*pl1*, 65% were found in IncI2 plasmids, and 43% of contigs carrying two copies of ISA*pl1* belonged to IncHI2 plasmids. Conversely, the common IncX4 plasmids carried only one transposon with two copies of ISA*pl1* and none with a single copy of the element.

We identified two extended plasmid backbone sequences that could be aligned. The first such alignment encompasses a shared sequence of 7,161bp between 108 plasmid backgrounds and has been previously referred to as ‘Type A’^13^. These sequences contain 54 sequences co-occurring with an IncI2 replicon, with 54 of unknown plasmid type, and encompass a large fraction of the genetic diversity found in the *mcr-1* transposon, although a large proportion (9/108) belonged to the dominant sequence type (i.e. B07_1_WFA61 in Figure 4). The second alignment is 34,761 bp long and is common to nine IncX4 plasmids and partly overlaps with a background previously defined as ‘Type D’^13^.

We applied BEAST to infer a timed phylogeny for each of these alignable regions after removal of SNPs showing evidence of recombination. For the IncI2 background we infer that a common ancestor to all 108 isolates existed in 2006 (1998-2010 95% CI relaxed exponential clock model) assuming a constant population size model (Figure S3). For the IncX4 backgrounds we dated the common ancestor of the eight isolates to 2011 (2010-2013 95% CI relaxed exponential clock model) assuming a constant population size model (Figure S6). Posterior density distributions of root dating under different population models are shown in Figures S2-S5. The difference in dating inferred for these two plasmid backgrounds and the recent date obtained for IncX4 highlight the dynamic nature of the integration of the *mcr-1* carrying transposon, even if in the IncX4 phylogeny isolates from East Asia and Europe and the Americas cluster together. The inferred mutation rates obtained for the IncI2 and IncX4 backgrounds consistently lie around 5-10x10^−5^ substitutions per site per year (Table S1).

### *Environmental distribution of the composite* mcr-1 *transposon*

It has been suggested that agricultural use of colistin, as has been widespread in China since the early 1980s^20^, caused the initial emergence and spread of *mcr-1*^24,25^. According to the evolutionary model in Figure 2, the ancestral mobilizable state is represented by the transposon carrying both its ISA*pll* elements. The transposon is thought to lose its capability for mobilization after the loss of both ISA*pll* elements^12^, although a single copy is reportedly sufficient to keep some ability to mobilize, with the upstream copy being functionally more important. Comparing human (n=108) and nonhuman (n=252) isolates, there were significantly more sequences with some trace of the insertion sequence ISA*pll* both upstream and downstream in non-human isolates (32/220 vs. 5/108, *χ^2^* test, *p*=0.033). This comparison held when only comparing agricultural isolates to human isolates (n=213) (28/213 vs. 5/108, *χ^2^* test, *p*=0.029). Furthermore, of the 42 isolates that had ISA*pll* fragments both upstream and downstream, the majority were from Asia (n=30) with only a quarter from Europe (n=10) (*χ^2^* test, *p*=0.12). This result is not driven by an over-representation of agricultural isolates from Asia in our dataset (*χ^2^* test, *p*=0.38)

## Discussion

We assembled a global dataset of 457 *mcr-1* positive sequenced isolates and could show that there was a single integration event of *mcr-1* into an ISA*pl1* composite transposon, followed by its subsequent spread between multiple genomic backgrounds. Our phylogenetic analyses suggest an age of insertion of *mcr-1* into the gene transposon shared across our isolates in the mid 2000s (2002-2008 95% HPD). We could identify the likely sequence of the ancestral transposon type and show the pattern of diversity supports a single mobilization with subsequent diversification during global spread.

Despite the limited number of whole genome sequences for samples before 2012, with the oldest sequence available from 2008 (Figure 1C), our estimate is consistent with the majority of available evidence from retrospective surveillance data^24^ which has found the presence of *mcr-1* in samples dating back to 2005 in Europe^26^. One retrospective study of Chinese isolates from 1970-2014 reported three *mcr-1* positive *E. coli* dating from the 1980s^27^, although *mcr-1* then did not reappear until 2004. If valid, this isolated observation provides evidence only for the presence of *mcr-1* as a gene rather than in mobilizable form, which would be consistent with an early emergence but a long dormancy before the formation of the composite transposon, subsequent mobilization, and global spread. We found that a constant population size model gave a better fit than an exponential model, suggesting the dramatic increase in reports of the presence of *mcr-1* across the past two years may not reflect a sudden global spread after its initial discovery^2^ and highlighting the difficulty of interpreting novel surveillance data from previously unknown resistance elements.

Our estimates of the age of spread of the representative IncI2 and IncX4 plasmid backgrounds are more recent, dating to around 2008 and 2013, respectively, but are both consistent with the age of the transposon mobilization event. We did not constrain the evolutionary rates in any of our phylogenetic analyses. It is thus encouraging that the different rates are highly consistent between the *mcr-1* transposon and the two plasmid backgrounds. While this points to high internal consistency between our estimates, we were surprisingly unable to find any previously published estimates for the evolutionary rate of bacterial plasmids.

The current distribution and observed genetic patterns are in line with a centre of origin in China. This is the place where we observe the highest proportion of isolates carrying intact or partial copies of the ISA*pl1* flanking elements. Transposon sequences carrying *ISA*pl1** elements were also overrepresented in environmental and agriculture isolates, relative to those collected from humans. This pattern is in line with agricultural settings acting as the source of *mcr-1* within bacteria isolated from humans^20^. The current global distribution has been achieved through multiple translocations, and is illustrated by the interspersed geographic origins in our phylogenetic reconstructions. A likely driver for the global spread is trade, in particular food animals^28^ and meat, although direct global movement by colonized or infected humans^19^ is also likely to have played a role in the current distribution.

The origin of *mcr-1* prior to its mobilization remains elusive. Despite an exhaustive search of sequence repositories, including the short-read archive, we found not a single *mcr-1* sequence outside the ISA*pl1* transposon background. ISA*pl1* was first identified in the pig pathogen *Actinobacillus pleuropneumoniae*^29^ suggesting that it may also have been an ancestral host for *mcr-1*, although to our knowledge no *mcr-1* positive *A. pleuropneumoniae* isolates have been described. The phosphoethanolamine transferase (*EptA)* from *Paenibacillus sophorae* has also been proposed as a possible candidate^30^. However, this seems most unlikely as *Paenibacili* are gram positive and are thus intrinsically resistant to polymixins^31^. Moreover, while the two sequences share functional similarities, this should be interpreted as a case of possible parallel evolution rather than direct filiation^31^. *Moraxella* has also been suggested as being the source of *mcr-1^32^*, following the identification of genes in *Moraxella* with limited homology to *mcr-1* (~60% nucleotide sequence identity). However, this sequence identity is too low for *Moraxella* to be considered as viable candidates for the origin of *mcr-1*. The search for the initial source of *mcr-1* remains open until a *mcr-1* sequence is identified outside of the ISA*pl1* sequence background.

We note that there are an increasing number of mobilized genes that can confer colistin resistance, with *mcr-2* reported less than a year after *mcr-1* was initially described^33^ and more recently the phylogenetically distant *mcr-3, mcr-4*, and *mcr-5* have also been described^34^^–^^36^. There appear to be commonalities between the mechanisms of the *mcr* genes, despite their different sequences and location near to different insertion sequences. For example, *mcr-2* has 76.7% nucleotide identity to *mcr-1* and was found in colistin-resistant isolates that did not contain *mcr-1*, and appeared to be mobilized on an IS1595 transposon^33^. Despite the different insertion sequences, intriguingly, this mobile element also contained a similar protein downstream of the *mcr* gene. Indeed, in *mcr-1, -2* and *-3*, the *mcr* gene has a downstream open reading frame encoding, respectively, a putative PAP2 protein^12^, a PAP2 membrane-associated lipid phosphatase^33^, and a diacylglycerol kinase^34^, all of which have transmembrane domains and are involved in the phosphatidic acid pathway^37,38^. While the PAP2-like orf in *mcr-1* has been shown to not be required for colistin resistance^39^, the presence of similar sequences downstream of other *mcr* genes implies some functional role, either in the formation of the mobile element and/or in its continued mobilization.

In summary, we assembled the largest dataset to date on *mcr-1* positive sequenced samples through our own sequencing efforts combined with an exhaustive search of *mcr-1* positive isolates sequenced and deposited on publicly available databases including unassembled datasets from the SRA. While this allowed us to obtain a truly global dataset of 457 *mcr-1* positive isolates covering 31 countries and five continents, we appreciate that the data is likely affected by complex sampling biases, with an overrepresentation of samples from places with active surveillance and well-funded research communities. Equally, although we took advantage of the most sophisticated bioinformatics and phylogenetic tools currently available, the complex “Russian doll” dynamics of the transposon, plasmids, and bacterial host limits our ability to reach strong inferences on some important aspects of the spread of *mcr-1*. Nevertheless, we believe our results highlight the potential for phylogenetic reconstruction of antimicrobial resistance elements at a global scale. We hope that future efforts relying on more sophisticated computational tools and even more extensive genetic sequence data will become part of the routine toolbox in infectious disease surveillance.

## Methods

### Compilation of genomic dataset

We blasted for *mcr-1* in all NCBI GenBank assemblies (as of 16th March 2017, *n*=90,759) using a 98% identity cut off. 195 records (0.21%; 121 assemblies, 73 complete plasmids, 1 complete chromosome) contained at least one contig with a full-length hit to *mcr-1* (1,626 bases). We only included samples with a single copy of *mcr-1*.

We searched a snapshot of all whole-genome sequenced bacterial raw read datasets in the SRA (December 2016), looking for samples containing mcr-1 by using a k-mer-index which we had previously constructed (k=31, software available at: https://github.com/phelimb/cbg). This snapshot consisted of 455,632 samples, of which 7,799 were excluded as they exceeded an arbitrary threshold of 10 million k-mers after error-cleaning with McCortex^40^. 184 datasets were found to contain at least 70% of the 31-mers in mcr-1. After removing duplicates (i.e. those with a draft assembly available) we could assemble contigs with *mcr-1* for 153 of these.

Our final dataset comprised 457 isolates from six genera across 31 different countries, ranging in date from 2008 to 2017. Where only a year was provided as the date of isolate collection the date was set to the midpoint of that year.

Whenever identified isolates did not comprise previously assembled genomes or complete plasmids, the raw fastq files were first inspected using FastQC and trimmed and filtered on a case by case basis. De-novo assembly was then conducted using Plasmid SPADES 3.10.0 using the --*careful* switch and otherwise default parameters ^41^. For those isolates sequenced using PacBio a different pipeline was employed. Correction, trimming and assembly of raw reads was performed using Canu^42^ and assembled reads were corrected and trimmed using the tool Circlator^43^. The quality of resultant assemblies was assessed using infoseq. In both cases the *mcr-1* carrying contigs in this final dataset were identified using blastn v2.2.31^44^.

We ran Plasmid Finder 1.3^45^ with 95% identity to identify plasmid replicons on the *mcr-1* carrying contigs. 177 contigs could be assigned a plasmid type using this method.

### Novel samples from China

We selected 110 *mcr-1* positive isolates from China for whole genome sequencing from a larger survey effort of both clinical and livestock isolates. 2,824 non-repetitive clinical isolates, including 1,637 *E. coli* and 1,187 *K. pneumoniae* were collected from 15 provinces of mainland China from 2011 to 2016. 72 isolates were resistant to polymyxin B, comprising 40 *E. coli* and four *K. pneumoniae* carrying *mcr-1*. Livestock samples were collected from four provinces of China in 2013 and 2016. One broiler farm of the Shandong province provided chicken anal swabs, liver, heart and wastewater isolated in 2013. In 2016, samples including faeces, wastewater, anal swabs, and internal organs of sick livestock were collected from swine farms, cattle farms and broiler farms in four provinces (Jilin, Shandong, Henan and Guangzhou). A total of 601 *E. coli* and 126 *K. pneumoniae* were isolated, of which 167 (137 *E. coli* and 30 *K. pneumoniae)* were resistant to polymyxin B. We detected *mcr-1* in 135 *E. coli* and two *K. pneumoniae*, as well as in eight *E. coli* isolated from environmental samples, which were collected from influents and effluents of four tertiary care teaching hospitals.

All of the isolates were sent to the microbiology laboratory of Peking University People’s Hospital and confirmed by routine biochemical tests, the Vitek system (bioMeriéux, Hazelwood, MO) and/or MALDI-TOF (Bruker Daltonics, Bremen, Germany). The minimal inhibitory concentrations (MICs) of polymyxin B was determined using the broth dilution method. The breakpoints of polymyxin B for Enterobacteriaceae were interpreted with the European Committee on Antimicrobial Susceptibility Testing (EUCAST, http://www.eucast.org/clinical_breakpoints) guidelines. Colistin-resistant isolates (MIC of ≥ 2 μg/ml) were screened for *mcr-1* by PCR and sequencing as described previously^46^.

### Identification and alignment of mcr-1 transposon

We searched for the *mcr-1* carrying transposon region across isolates by blasting for its major components: *ISA*pl1* (Actinobacillus pleuropneumoniae* reference sequence: EF407820), *mcr-1* (from *E. coli* plasmid pHNSHP45: KP347127.1), and short sequences representing the sequences immediately upstream and downstream of *mcr-1* (from KP347127.1) using blastn-short. Contiguous sequences containing *mcr-1* were aligned using Clustal Omega^47^ and then manually curated and amended using jalview ^48^, resulting in a 3,679bp alignment containing the common ~2,600bp identified by Snesrud and colleagues^12^. The downstream copy of ISA*pl1* was more often fragmented or inverted. 28 isolates which were all assemblies from the same study in Vietnam had a ~1.7kb insertion downstream of *mcr-1* (Figure 3e) before the downstream ISA*pl1* element.

### Phylogenetic analyses

For constructing the transposon phylogeny, we excluded the downstream ISA*pl1* and the insertion sequence observed in a small number of samples, as well as regions identified as having signals of recombination by ClonalFrameML^49^, resulting in a 3,522bp alignment. We removed two homoplastic sites (requiring >1 change on the phylogeny), before constructing a maximum parsimony neighbor-joining tree based on the Hamming distance between sequences. We calculated branch lengths using nonnegative least squares with nnls.phylo in phangorn v2.2.0^50^. Phylogenies were visualized with ggtree v1.8.1^51^.

### Phylogenetic dating

Given recombination can conceal the clonal phylogenetic signal we also applied ClonalFrameML^49^ to identify regions of high recombination in a subset of IncI2 and IncX4 plasmid background alignments. Where recombination hotspots were identified, they were removed from the alignment. In the IncI2 alignment this resulted in removing 1,281 positions. No regions of high recombination were detected in the IncX4 alignment. We applied root-to-tip correlations to test for a temporal signal in the data using TempEST^52^. There was a significantly positive slope for all three alignments (Fig. S7-S9).

We applied BEAUTi and BEAST v2.4.7^53,54^ to estimate a timed phylogeny from an alignment of IncI2 plasmids (7,161 sites, 110 isolates) and IncX4 plasmids (34,761 sites, 8 isolates). Sequences were annotated using their known sampling times expressed in years. For both plasmid alignments, the HKY substitution model was selected based on evaluation of all possible substitution models in bModelTest. Beast analyses were then applied under both a coalescent population model (the coalescent Bayesian skyline implementation) and an exponential growth model (Coalescent Exponential population implementation). Additionally, a strict clock, with a lognormal prior, and a relaxed clock (both lognormal and exponential) were tested. MCMC was run for 50,000,000 iterations sampling every 2000 steps and convergence was checked by inspecting the effective sample sizes (ESS) and parameter value traces in the software Tracer v1.6.0. Analyses were repeated three times to ensure consistency between the obtained posterior distributions. Posterior trees for the best fitting model were combined in TreeAnnotator after a 10% burn-in to provide an annotated Maximum Clade Credibility (MCC) tree. MCC trees were plotted using ggtree^51^ for both backgrounds: IncI2 (Figure S10) and IncX4 (Figure S11). The model fit across analyses was compared using the Akaike’s information criteria model (AICM) through 100 boot-strap resamples as described in Baele and colleagues^55^ and implemented in Tracer v1.6 (Table S2).

Phylogenetic dating on the transposon was performed using an alignment of 364 isolates, which included only those with information on isolation date, across 3,522 sites. As before Beast analyses were applied under both a coalescent population model (coalescent Bayesian skyline implementation) and an exponential growth model (coalescent exponential population implementation). Additionally, a strict clock, with a lognormal prior, and a relaxed clock (both lognormal and exponential) were tested. Analyses were run under a HKY substitution model for 600 million iterations sampling every 5,000 steps. Only analyses using a strict clock model reached convergence after 600 million iterations. The resultant set of trees were thinned by sampling every 10 trees and excluding a 10% burn-in and combined using TreeAnnotator to produce a MCC tree. MCC trees were plotted using ggTree^48^. As before the model fit was evaluated using AICM’s implemented in Tracer v1.6.

### Environmental distribution

For the purpose of testing the distribution of sequences containing some trace of *ISA*pll*, we* classed isolates into broad categories as either environmental (n=39; bird, cat, dog, fly, food, penguin, reptile, vegetables), agricultural (n=213; chicken, cow, pig, poultry feed, sheep, turkey), or human (n=108).

## Acknowledgments

L.v.D., X.D., H.W., T.S., L.A.W. and FB acknowledge financial support from the Newton Trust UK-China NSFC initiative (grants MR/P007597/1 and 81661138006). F.B. additionally acknowledges support from the BBSRC GCRF scheme. L.S. was supported by a PhD scholarship from EPSRC (EP/F500351/1). P.B. is funded by Wellcome Trust on a “Genomic Medicine and Statistics DPhil” grant. A.R. was co-funded by the European Union: European regional development fund (ERDF), by the Conseil Régional de La Réunion and by the Centre de Coopération internationale en Recherche agronomique pour le Développement (CIRAD). H.W. was supported by National Natural Science Foundation of China (81625014). L.A.W. is supported by a Dorothy Hodgkin Fellowship funded by the Royal Society (Grant Number DH140195) and a Sir Henry Dale Fellowship jointly funded by the Wellcome Trust and the Royal Society (Grant Number 109385/Z/15/Z). The funders had no role in study design, data collection and interpretation, or the decision to submit the work for publication.

## Data access

The 110 *mcr-1* positive Illumina sequencing data have been submitted to the Short Read Archive under Bioproject: <u>PRJNA408214</u>, Accession: SRP118547. The data will be released upon peer-reviewed publication or on the 2018-9-21.

## Contributions

H. W., and F.B. conceived the project and designed the experiments. R.W., Q.W., X.W., L.J. Q.Z., Y.L and H.W collected samples. R.W., Q.W., X.W., L.J., Q.Z., and Y.L performed microbial identification, antimicrobial susceptibility testing, screening for *mcr-1*, and DNA extraction for WGS. R.W. L.V.D., L.S. assembled the new sequence data. L.v.D. and L.S. curated the global dataset and performed the computational analyses. T.D.S. advised on functional aspects of colistin resistance. A.R, L.A.W. and X.D. helped with the phylogenetic reconstructions. P.B. and Z.I. performed the search for *mcr-1* positive samples on the Short Read Archive. H.W. takes responsibility for the accuracy and availability of the epidemiological and raw sequence data, and F.B. for all bioinformatics and computational methods and results. L.v.D., L.S. and F.B. wrote the paper with contributions from X.D. and H.W. All authors read and commented on successive drafts and all approved the content of the final version. Z.I. was funded by a Sir Henry Dale Fellowship jointly funded by the Wellcome Trust and the Royal Society (Grant Number 102541/Z/13/Z)

